# Defining Mitochondrial Cristae Morphology Changes Induced by Aging in Brown Adipose Tissue

**DOI:** 10.1101/2023.05.12.540609

**Authors:** Amber Crabtree, Kit Neikirk, Andrea G. Marshall, Larry Vang, Aaron J. Whiteside, Qiana Williams, Christopher T. Altamura, Trinity Celeste Owens, Dominique Stephens, Bryanna Shao, Alice Koh, Mason Killion, Edgar Garza Lopez, Jacob Lam, Ben Rodriguez, Margaret Mungai, Jade Stanley, E. Danielle Dean, Ho-Jin Koh, Jennifer A. Gaddy, Estevão Scudese, Mariya Sweetwyne, Jamaine Davis, Elma Zaganjor, Sandra A. Murray, Prasanna Katti, Steven M. Damo, Zer Vue, Antentor Hinton

**Affiliations:** Department of Molecular Physiology and Biophysics, Vanderbilt University, Nashville, TN, 37232, USA; Department of Internal Medicine, University of Iowa, Iowa City, IA, 52242, USA; Division of Diabetes, Endocrinology, and Metabolism, Department of Medicine, Vanderbilt University Medical Center, TN, 37232, USA; Department of Medicine, Vanderbilt University Medical Center, Nashville, TN, 37232, USA; Department of Biological Sciences, Tennessee State University, Nashville, TN 37209; Tennessee Valley Healthcare Systems, U.S. Department of Veterans Affairs, Nashville, TN, 37232, USA; Laboratory of Biosciences of Human Motricity (LABIMH) of the Federal University of State of Rio de Janeiro (UNIRIO), Rio de Janeiro, Brazil; Sport Sciences and Exercise Laboratory (LaCEE), Catholic University of Petrópolis (UCP), Brazil; Department of Laboratory Medicine and Pathology, University of Washington, Seattle, WA, 98195, USA; Department of Biochemistry, Cancer Biology, Neuroscience, Pharmacology, Meharry Medical College, Nashville, TN 37208 USA; Department of Cell Biology, University of Pittsburgh; Pittsburgh, PA, 15261 USA; National Heart, Lung and Blood Institute, National Institutes of Health, 9000 Rockville Pike, Bethesda, MD 20892, USA; Department of Life and Physical Sciences, Fisk University, Nashville, TN, 37208, USA; Center for Structural Biology, Vanderbilt University, Nashville, TN, 37232, USA

## Abstract

Mitochondria are required for energy production and even give brown adipose tissue (BAT) its characteristic color due to their high iron content and abundance. The physiological function and bioenergetic capacity of mitochondria are connected to the structure, folding, and organization of its inner-membrane cristae. During the aging process, mitochondrial dysfunction is observed, and the regulatory balance of mitochondrial dynamics is often disrupted, leading to increased mitochondrial fragmentation in aging cells. Therefore, we hypothesized that significant morphological changes in BAT mitochondria and cristae would be present with aging. We developed a quantitative three-dimensional (3D) electron microscopy approach to map cristae network organization in mouse BAT to test this hypothesis. Using this methodology, we investigated the 3D morphology of mitochondrial cristae in adult (3-month) and aged (2-year) murine BAT tissue via serial block face-scanning electron microscopy (SBF-SEM) and 3D reconstruction software for manual segmentation, analysis, and quantification. Upon investigation, we found increases in mitochondrial volume, surface area, and complexity and decreased sphericity in aged BAT, alongside significant decreases in cristae volume, area, perimeter, and score. Overall, these data define the nature of the mitochondrial structure in murine BAT across aging.

Graphical Abstract:
Overview of serial block facing-scanning electron microscopy (SBF-SEM) workflow, data segmentation, and 3D analysis of mitochondria using Amira software for murine interscapular BAT.

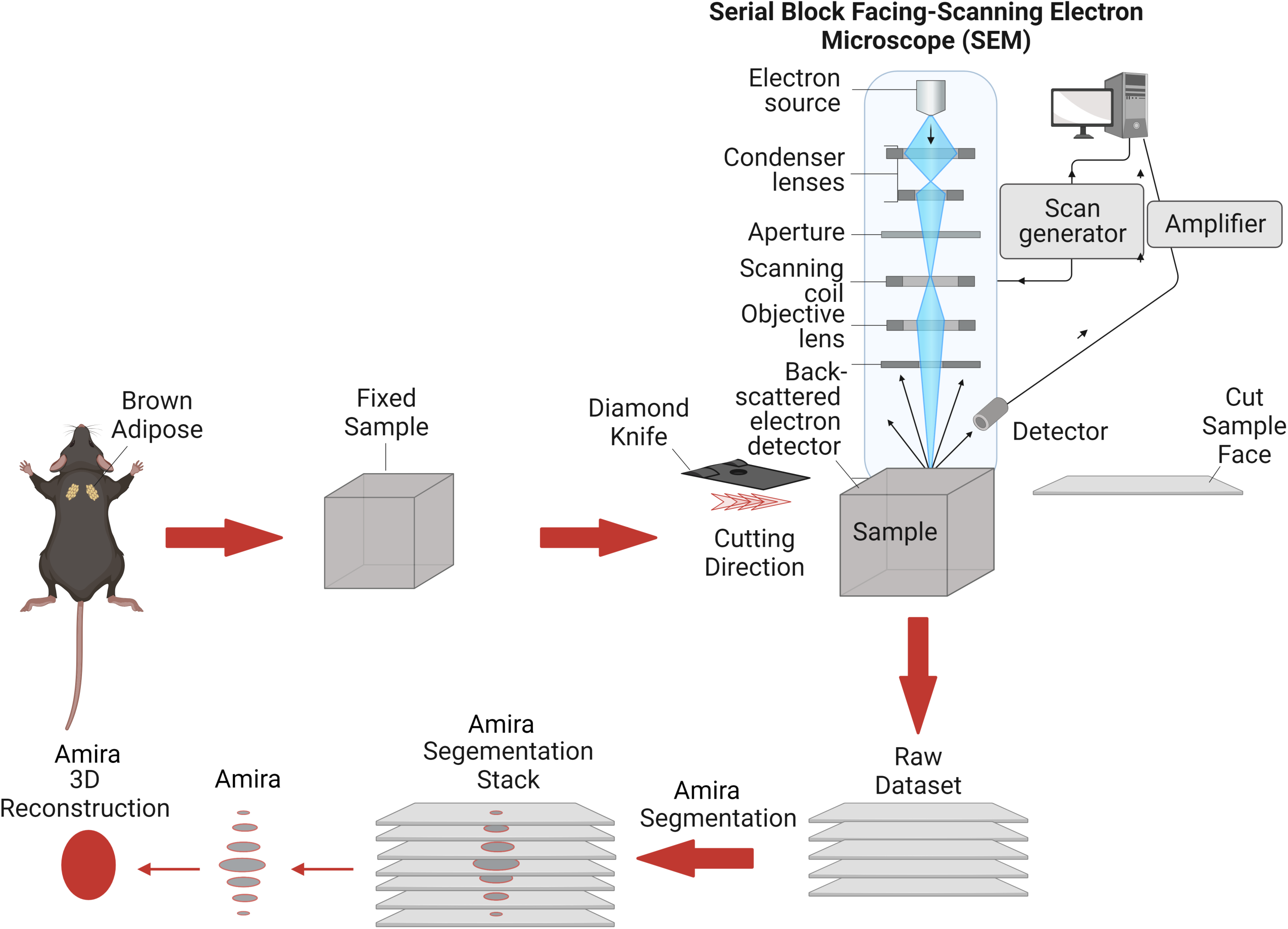

## INTRODUCTION

Mitochondria are complex cellular organelles that serve various physiological roles, including maintenance of Ca^2+^ homeostasis, initiation of apoptosis, and cellular energy production ^[1–3]^. With such vital cellular roles, mitochondria must be dynamic to meet the varying energetic demands of the cell ^[1, 4–6]^. The ultrastructure and morphology of mitochondria is tightly associated with their functional capacity ^[7]^. Therefore, it is no surprise that mitochondrial morphology varies considerably between tissue types and the metabolic health of that tissue ^[4, 8]^. The invaginations of the inner mitochondrial membrane (IMM), known as cristae, substantially increase the surface area for the oxidative phosphorylation machinery to reside, optimizing energetic capacity ^[7]^. Therefore, maintaining the structural integrity and spatial arrangement is essential for maintaining bioenergetic homeostasis and ensuring that energetic demands can be met ^[9]^.

Normal mitochondrial dynamics consist of well-orchestrated, balanced cycles of fusion and fission that allow for content mixing and quality control to maintain a healthy network ^[6,10,11]^. While it was previously believed that mitochondrial dynamics only occurred in response to changes in the cellular environment, it is now known that mitochondrial dynamics are required for maintaining healthy mitochondrial networks and that cristae undergo similar fusion-like and fission-like events ^[7, 12]^. Despite the identification of key regulators for cristae morphology, our understanding of the dynamics of cristae reorganization in physiological and disease contexts remains limited ^[13, 14]^.

Mitochondrial oxidative stress and dysfunction are often associated with the pathophysiology of diseases and the aging process, which can lead to the accumulation of mitochondrial DNA (mtDNA) mutations as well as the generation of reactive oxygen species ^[15–, 17]^. The mitochondrial ultrastructure changes that occur with the aging process are only now possible to explore with the development of 3D reconstruction techniques. For example, studies have shown that in murine brain, the hippocampal, somatic, dendritic, and axonal mitochondria have differential baseline phenotypes and responses to aging ^[18]^. Recently, 3D reconstruction studies revealed that in aged murine heart, the mitochondria arranged in a less ordered manner than their younger counterparts ^[19]^. These findings establish the applicability of 3D reconstruction to study mitochondrial phenotypes and cellular organization across aging.

The number of mitochondria present, along with their form and function, are highly tissue-dependent ^[8, 20, 21]^. Brown adipose tissue (BAT) is rich in mitochondria, which are integral in the process of thermogenesis ^[22]^. Brown adipocytes produce heat primarily through the uncoupling of the cristae’s proton gradient, which is facilitated through uncoupling protein 1 (UCP1) ^[23]^. In both rodent and human models, a decrease in BAT thermogenic function has been observed with advanced aging and has been associated with the development of metabolic disorders, including obesity and diabetes. Studies suggest that mitochondrial dysfunction occurs through decreased expression of UCP1, and the functional age-related decline in BAT has been associated with mitochondrial dysfunction in several studies ^[23–25]^. In oxidative-stress-induced models of aging, BAT mitochondria showed decreased expression of UCP1, increased autophagy, and decreased overall size ^[26]^. Yet, to our knowledge, no studies have elucidated changes in the 3D morphology of mitochondria and cristae, as well as their 3D spatial distribution in BAT.

In this study, serial block face-scanning electron microscopy (SBF-SEM) and 3D reconstruction software were utilized to examine the 3D architecture of both mitochondria and cristae in BAT from adult and aged mice. Validated quantification methods ^[27]^ were used to establish a quantitative 3D approach to map mitochondrial networks and cristae. Here, we show that mitochondrial 3D volume, perimeter, 3D area, and complexity index all increased, while mitochondrial sphericity decreased, with aging in murine BAT. Conversely, mitochondrial cristate showed decreases in 3D volume, perimeter, 3D area, and cristae score with aging.

## METHODS

### Mice Care Procedure

Male C57BL/6J mice were housed at 22LJ C with a 12-h light, 12-h dark cycle accompanied by free access to water and standard chow following birth. Mice were cared for as previously discussed ^[28]^ with all protocols approved by the University of Iowa Animal Care and Use Committee (IACUC).

### SBF-SEM Sample Preparation

Interscapular BAT was excised from 3-month and 2-year aged mice and fixed in 2% glutaraldehyde in 0.1M cacodylate buffer and processed using a heavy metal protocol. BAT samples were then immersed in 3% potassium ferrocyanide, followed by 2% osmium tetroxide for 1 hour each at 4LJ C following deionized H_2_O (diH_2_O) washes. Samples were washed again in diH_2_O and immersed in filtered 0.1% thiocarbohydrazide for 20 minutes, followed by diH_2_O washing and subsequent immersion in 2% osmium tetroxide for 30 minutes. Samples were incubated overnight in 1% uranyl acetate at 4LJ C. The following day, samples were incubated in 0.6% lead aspartate for 30 minutes at 60LJ C prior to an ethanol graded series dehydration. BAT tissue samples were then infiltrated with epoxy Taab 812 hard resin prior to immersion in fresh resin and polymerization at 60LJ C for 36-48 hours. The resultant resin blocks were sectioned for transmission electron microscopy (TEM) to identify regions of interest. Samples were then trimmed, glued to an SEM stub, and then placed into a FEI/Thermo Scientific Volumescope 2 SEM. Between 300-400 thin serial sections of 0.09 μm per sample block were obtained and collected. The resultant micrograph blocks were then aligned and manually segmented and reconstructed in 3D using Thermo Scientific Amira Software (Waltham, MA, USA) ^[27, 29, 30]^.

### Mitochondrial and Cristae Ultrastructure Calculations and Measurements

Following manual segmentation of mitochondria and cristae in the regions of interest (ROIs), label analyses were performed on each segmented structure using Amira ^[27]^. The SBF-SEM data was acquired from at least three independent experiments to perform blinded-3D structural reconstruction from murine BAT. Manual segmentation of sequential orthoslices was then performed to obtain 300-400 slices of which, 50-100 serial sections were chosen for each 3D reconstruction. Serial sections had approximately equal z-direction intervals and werestacked, aligned, and visualized using Amira software to make videos and quantify volumetric structures. The algorithms for measurements were entered manually (all measurements in the main text) for those not already in the system, and a total of 300 mitochondria from three mice were collected for each age cohort (600 mitochondria/6 mice total).

### Structure Quantifications and Statistic Analyses

All data obtained from label analyses and manual measurements were statistically analyzed by Student’s t-test, or the non-parametric equivalent where applicable, using GraphPad Prism (San Diego, California, USA). All data considered are biological replicates and dots represent individual data points unless otherwise noted. In some cases, for presentation, certain outliers may not be displayed, but all outliers are considered for statistical analysis. Graphs are shown as means with bars representing the standard error of mean. A minimum threshold of p < 0.05 indicated a significant difference. Higher degrees of statistical significance (i.e., **, ***, ****) were defined as p < 0.01, p < 0.001, and p < 0.0001, respectively.

## RESULTS

### Mitochondrial and Matrix Volumes Significantly Increase with Aging In BAT

BAT biopsies were collected from adult (3-months-old) and aged (2-years-old) mice and imaged using SBF-SEM. With resolutions of 10 nm for the x- and y-planes and 50 nm for the z-plane, SBF-SEM allows for 3D reconstruction of organelles, providing a high spatial resolution that is unattainable using 2D techniques ^[31]^. To quantify mitochondrial changes across aging, we examined approximately 300 mitochondria from three separate regions of interest per age group were selected from the male mice (n=3) (Figure 1A). Stacks of ∼50 images of 50-µm block ortho slices (Figure 1B) were manually traced at transverse intervals (Figure 1C). This enabled the generation of 3D reconstructions of each mitochondrion (Figure 1D).

**Figure 1:**
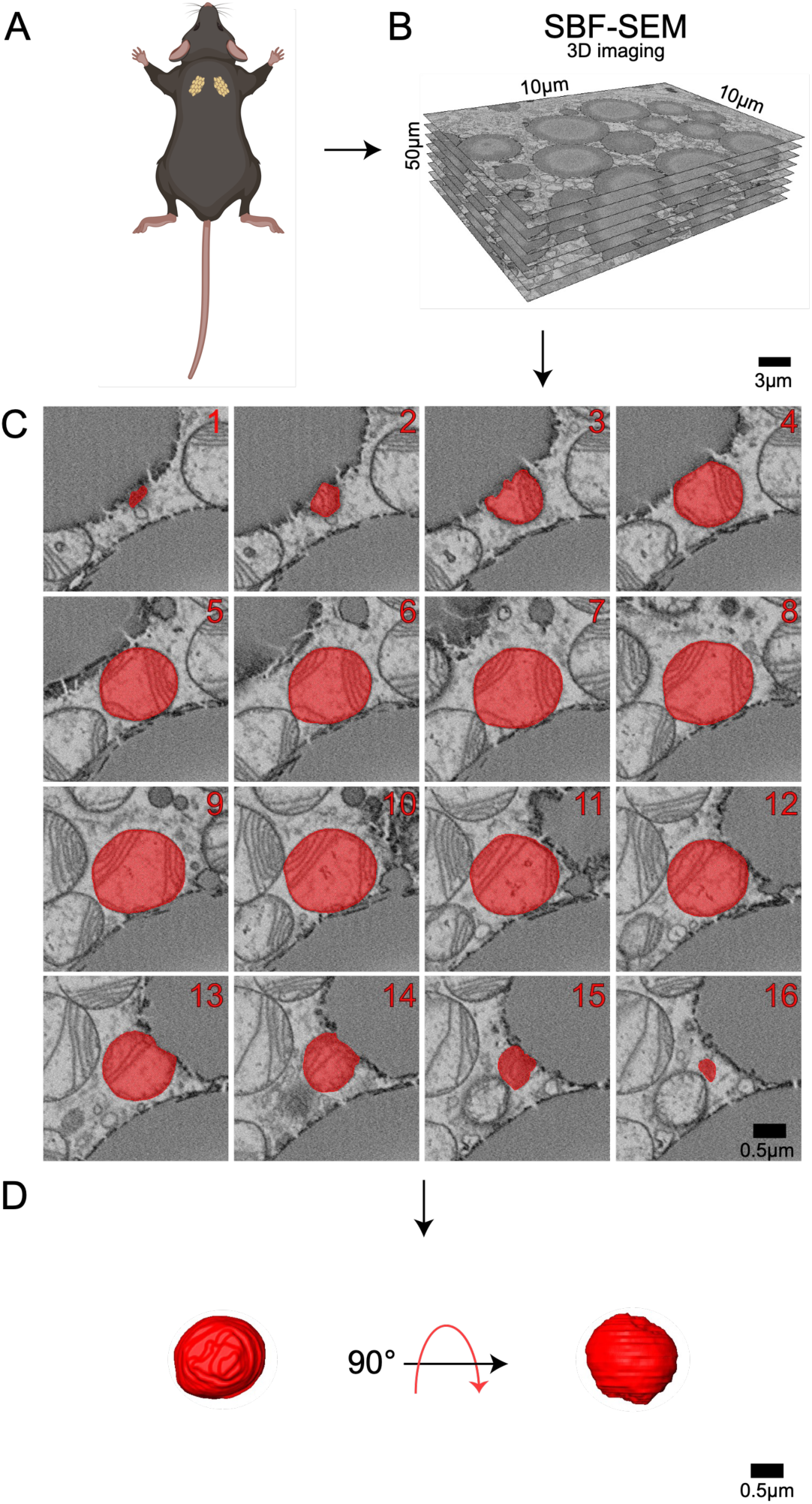
Detailed 3D reconstruction workflow for mitochondria using serial block face-scanning electron microscopy (SBF-SEM). (A) Interscapular brown adipose tissue (BAT) was carefully excised from male C57BL/6J mice. (B) High-resolution 10 µm by 10 µm orthoslices were generated by imaging consecutive ultrathin sections of the sample. These orthoslices were then accurately aligned and stacked to produce a comprehensive 3D volume of the tissue. (C) SBF-SEM images were processed to create representative orthoslices, which provided valuable information on the internal structure of the mitochondria. (D) A detailed 3D reconstruction of mitochondria was produced based on the orthoslices, allowing for in-depth analysis of mitochondrial morphology and spatial organization.

BAT mitochondria are typically abundant, large, and round in young, healthy tissue ^[32]^. Consistent with this, we observed an abundance of mitochondria that were spherical in structure in the adult (aged 3-months) group, which is believed to be equivalent to approximately 20-years in the human lifespan. In comparison, the 2-year aged samples are believed to be equivalent to approximately 70 years of age in humans ^[33]^. When comparing the 3-month to 2-year ages, significant increases in perimeter, 3D area (or surface area), and volume were observed (Figure 2). When comparing the mitochondrial quantifications from each mouse, they exhibited minimal inter-group heterogeneity but consistent intra-group variability with increased heterogeneity in aged samples (SFigure 1).

**Figure 2:**
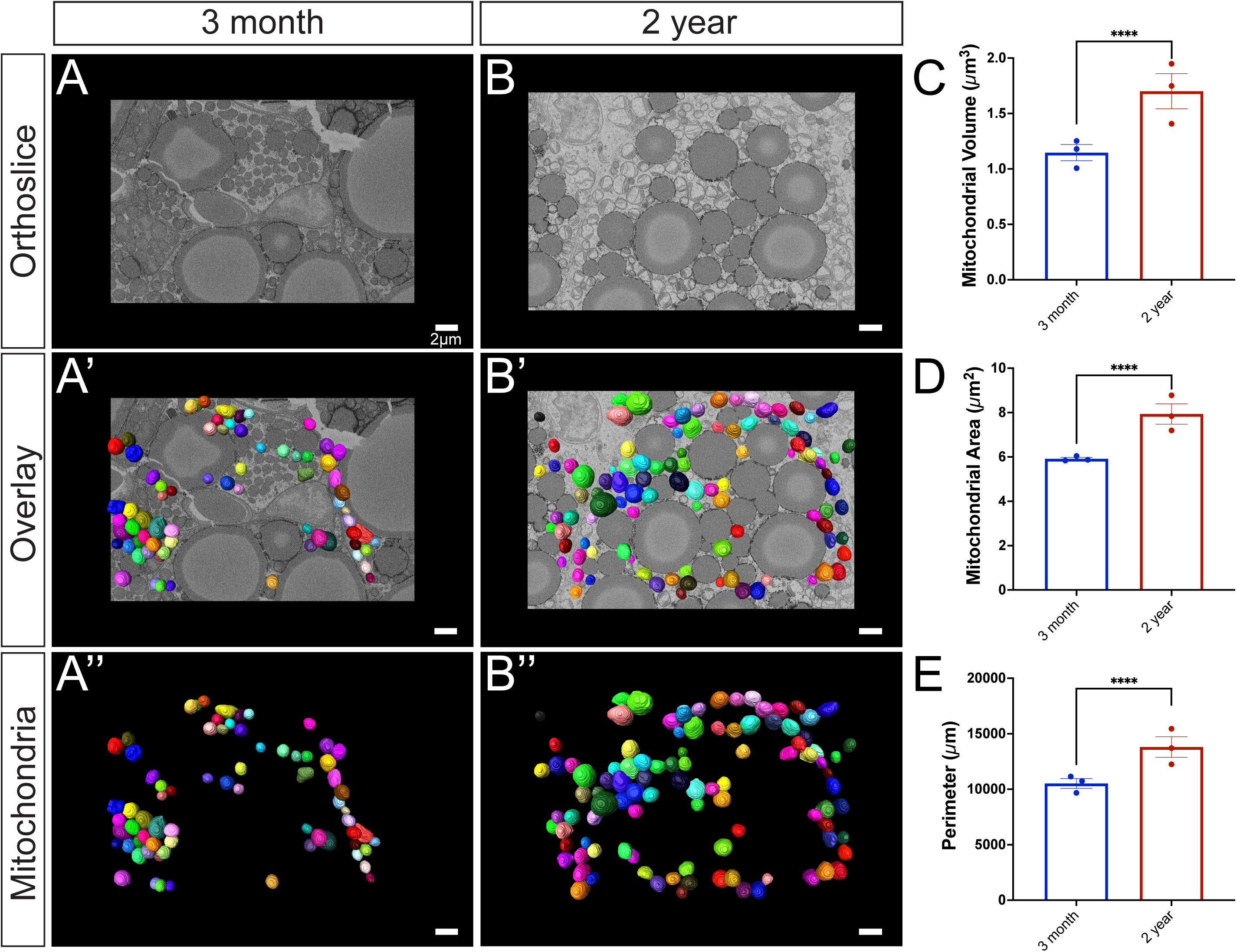
Comprehensive analysis of mitochondrial morphology changes during the aging process. (A) 3-month-old murine BAT: a representative orthoslice showcases the intricate internal mitochondrial structure; (A’) 3D reconstruction of mitochondria overlaid on the orthoslice illustrates the spatial distribution of mitochondria within the tissue;(A’’) Isolated 3D reconstruction with orthoslice removed, highlighting the individual mitochondrial structure. (B) 2-year-old murine BAT: a representative orthoslice depicts the aged mitochondrial structure; (B’) a 3D reconstruction of mitochondria overlaid on the orthoslice demonstrates the spatial organization of aged mitochondria; (B’’) Isolated 3D reconstruction with the orthoslice removed the individual aged mitochondrial structure. (C) Quantitative comparison of 3D mitochondrial volume, (D) surface area, and (E) perimeter between 3-month and 2-year samples to assess morphological differences, revealing age-related changes in size and shape of the mitochondria. Unpaired t-test was used to evaluate changes, with a variable sample number (n approximately 300), with each individual mice’s average represented by black dots. p < 0.05, p < 0.01, p < 0.001, and p < 0.0001 are indicated by *, **, ***, and ****, respectively.

### BAT Aging Significantly Increases Mitochondrial Complexity

Brown adipocytes are an integral part of BAT and are generally classified as either being high-thermogenic or low-thermogenic. High thermogenic brown adipocytes are characterized by smaller lipid size, mitochondria that are round in shape, and a high basal respiration rate ^[34]^. Low thermogenic brown adipocytes, on the other hand, are characterized by larger lipids, oval-shaped mitochondria, and a low basal respiration rate. Interestingly, a reduced thermogenic capacity is associated with aging and obesity ^[34]^. Thus, characterizing the shape of mitochondria may have important implications for their overall functional state.

Viewing the overall complexity of mitochondria can provide relative insights into the complexity of the mitochondrial networks they form as seen in skeletal and cardiac tissue ^[20]^. The sphericity of mitochondria, or the closeness something is in shape to a perfect sphere, shows a significant decrease across aging (Figure 3C and S1E). Of note, high-thermogenic BAT mitochondria are round whereas low-thermogenic mitochondria are considered oval in shape ^[34]^. The mitochondrial complexity index (MCI) is an index that correlates with the complexity of the mitochondrial shape, such as surface area and branching relative to volume (Surface Area^3^ /16 π^2^volume^2^). As an analogous measure for sphericity (calculated as 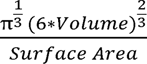), it remains consistent for mitochondria with the same shape but different volumes, as demonstrated by 3D model simulations ^[35]^. The MCI showed increases in median mitochondrial complexity across aging (Figure 3D and S1D). To visualize how mitochondrial complexity is changed across mitochondrial volume, we performed a technique known as mito-otyping, which organizes mitochondria based on their volume for each aging point (Figure 3E). Importantly, using this technique allowed us to simultaneously visualize changes in both complexity and volume across our aging model at each volume point analyzed.

**Figure 3:**
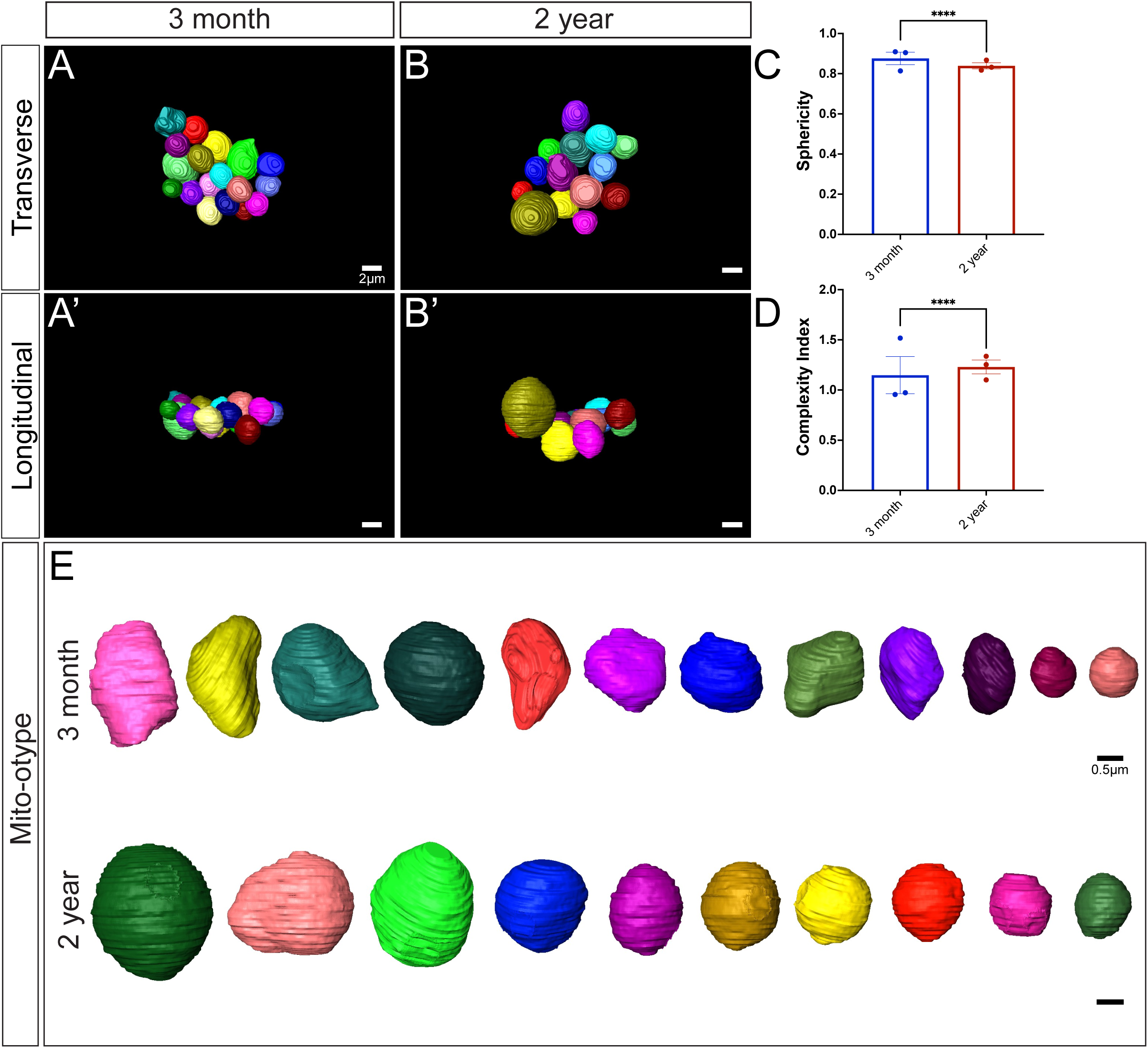
In-depth analysis of age-related changes in mitochondrial complexity. (A) 3-month-old murine BAT: 3D reconstruction of mitochondria overlaid from the transverse perspective (A’) a longitudinal perspective of the 3D reconstruction reveals additional details about the spatial organization of the mitochondria. (B) 2-year-old murine BAT: reconstruction of mitochondria overlaid on the orthoslice presents an overview of the aged mitochondrial network from transverse and (B’) a longitudinal perspective of the 3D reconstruction displays the spatial organization of the aged mitochondria. (C) Quantitative comparison of 3D sphericity and (D) complexity index; (E) Mito-otyping for mitochondrial morphology in 3-month and 2-year samples based on volume, providing insights into the age-related alterations in mitochondrial organization, shape, and structural complexity. Unpaired t-test was used to evaluate changes, with a variable sample number (n approximately 300), with each individual mice’s average represented by black dots. p < 0.05, p < 0.01, p < 0.001, and p < 0.0001 are indicated by *, **, ***, and ****, respectively.

### Aged BAT Mitochondria Show Decreased Cristae Volume, Area, Perimeter, and Score

Mitochondrial morphology is tightly associated with mitochondrial function, making it an important component that is often unexplored. Alterations in cristae morphology and matrix volume have the potential to provide important quantitative changes in BAT aging. Although there are morphological restraints, increased cristae volume generally correlates with an increased ATP potential ^[36]^. Notably, inflammation of BAT has been demonstrated to cause loss of cristae structural integrity, possibly related to impaired UCP1 activation, which is also observed ^[37]^. Therefore, we sought to quantify cristae morphological and size changes across aging. Using the workflow of SBF-SEM-based 3D reconstruction for BAT cristae, BAT was obtained from the same regions of interest on the mice (Figure 4A), and 10 µm by 10 µm orthoslices were overlaid for 3D reconstruction (Figure 4B). Representative orthoslices from the SBF-SEM sectioning were used to generate a detailed 3D reconstruction of cristae (Figure 4C-D).

**Figure 4:**
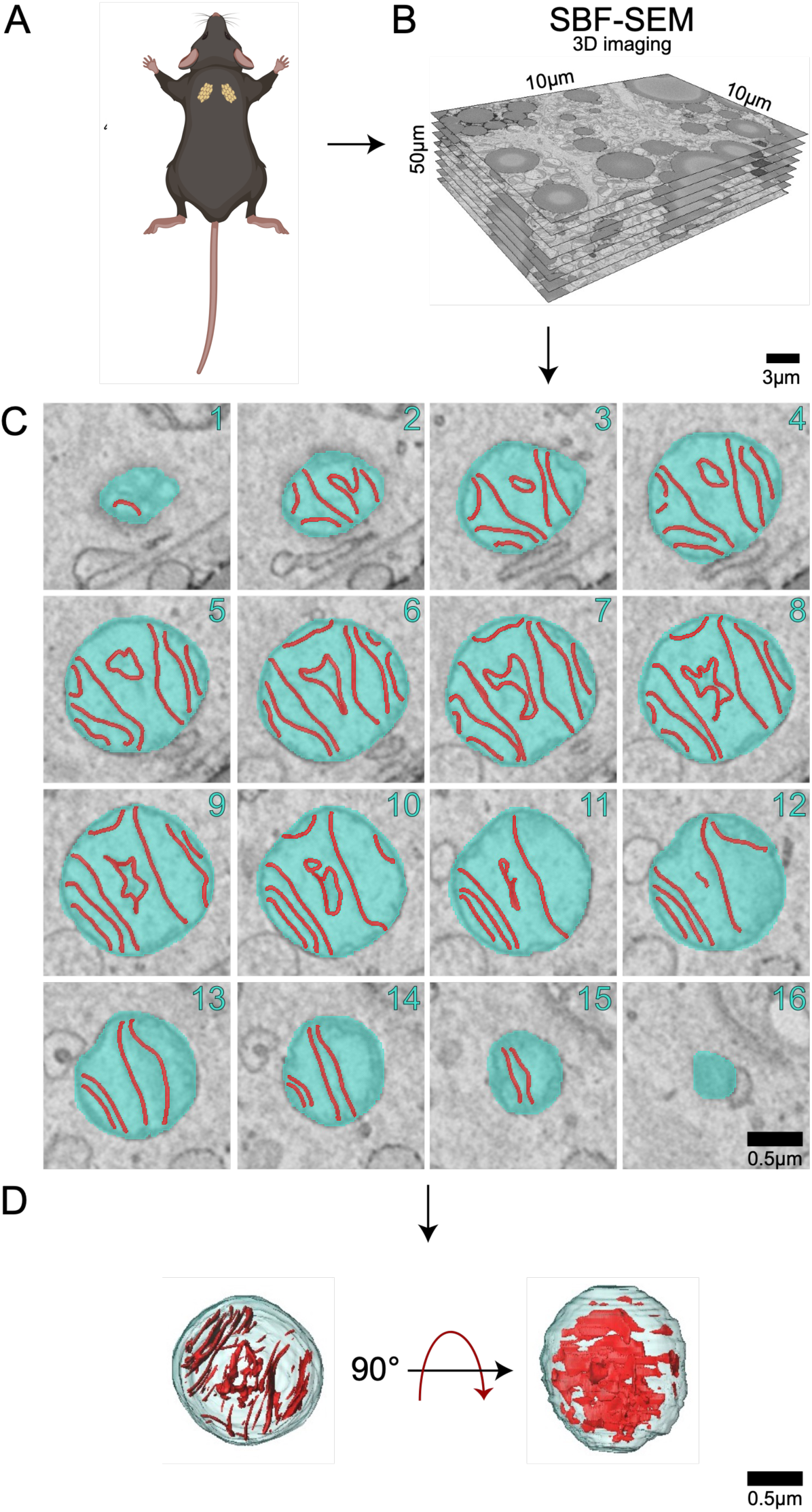
Workflow of serial block face-scanning electron microscopy-based 3D reconstruction for cristae. (A) Brown adipose tissue was excised from 3-month and 2-year aged male C57BL/6J mice. (B) 10 µm by 10 µm orthoslices were overlaid for 3D reconstruction. (C) Representative orthoslices from SBF-SEM slices to make (D) cristae 3D reconstruction.

Cristae and matrix morphology changes across aging indicate that cristae in the 2-year BAT are more heterogenous in size, but are generally smaller and more complex than those in the 3-month samples (Figure 5A-E). Interestingly, while the mitochondrial outer membrane parameters increased overall, we observe the opposite here, where cristae volume, surface area, and perimeter exhibit a significant decrease when normalized by mitochondrial volume. Despite increased heterogeneity and larger outliers (Supplemental Figure 2), this data indicates cristae density is reduced across aging in BAT when normalized by mitochondrial volume.

**Figure 5:**
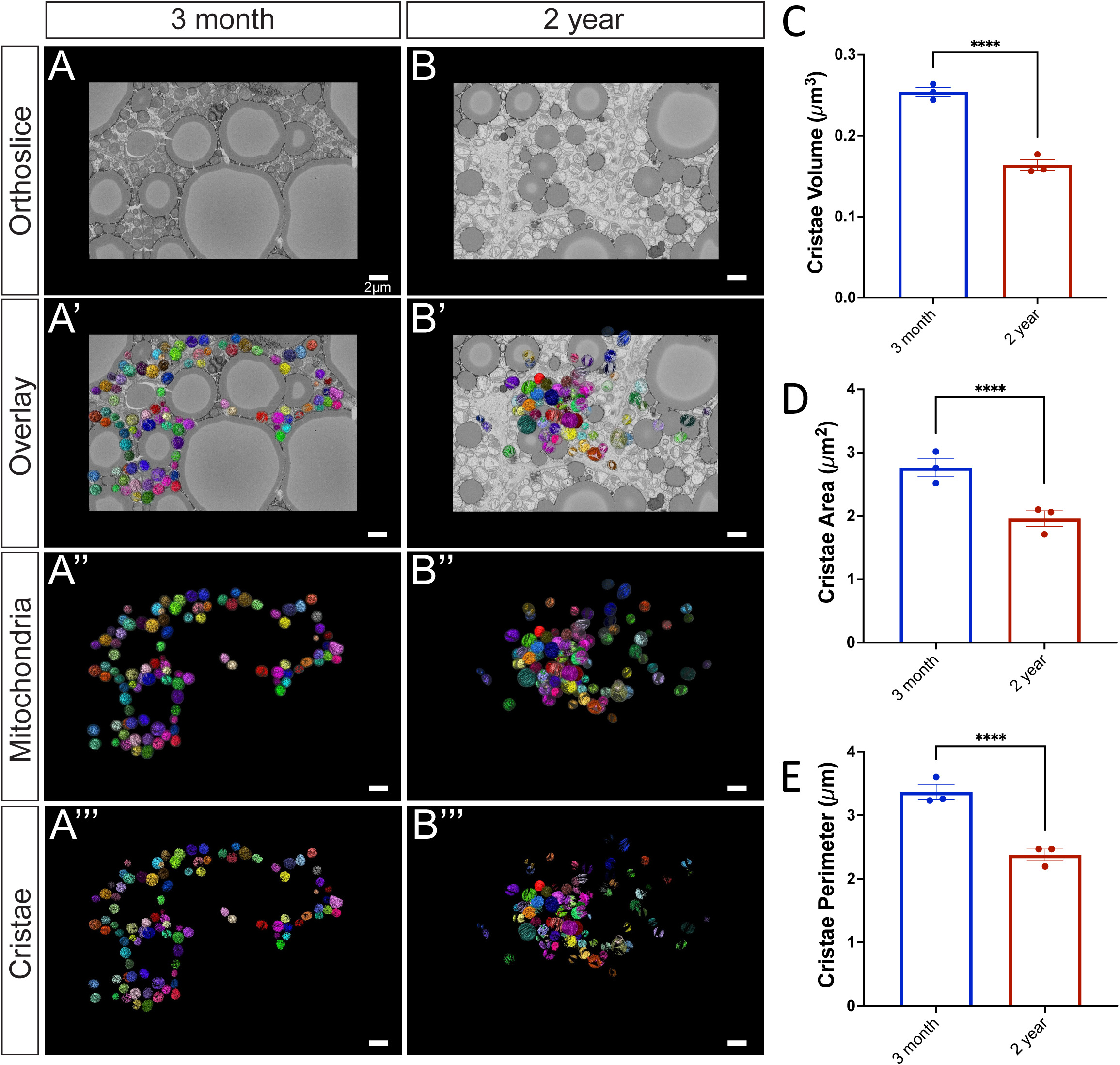
Cristae morphology across aging. (A) 3-month murine BAT representative orthoslice; (A’) 3D reconstruction of mitochondria matrix and cristae overlaid on orthoslice; (A’’) 3D reconstruction isolated mitochondrial matrix and cristae; (A’’’) 3D reconstructions of cristae alone. (B) 2-year murine BAT representative orthoslice; (B’) 3D reconstruction of mitochondria overlaid on orthoslice; (B’’) 3D reconstruction isolated mitochondrial matrix and cristae; (B’’’) 3D reconstructions of cristae alone. (C) Comparisons of cristae 3D volume, (D) 3D area, and (E) 3D perimeter compared in 3-month and 2-year samples. Unpaired t-test was used to evaluate changes, with a variable sample number (n approximately 300), with each individual mice’s average represented by black dots. p < 0.05, p < 0.01, p < 0.001, and p < 0.0001 are indicated by *, **, ***, and ****, respectively.

To better understand cristae changes across aging, we utilized the cristae score, which is a qualitative measurement assigned to cristae with a scale ranging from 1-4: a minimum score of 1 indicating minimal normal cristae, a score of 2 indicating less than 50% normal cristae, a score of 3 indicating 50-75% normal cristae, and a maximum score of 4 indicating normal cristae ^[28]^ (Figure 6A). While previously applied in transmission electron microscopy (EM), to our knowledge, we are the first to apply this scoring system to 3D EM. To eliminate the risk of unconscious bias, independent blinded cristae score analyses were performed. For the 2-year BAT samples, we noticed that both the frequency and percentage of cristae were ranked as a 1 or 2, while there were significantly less cristae ranked as a 3 (Figure 6B-C). Collectively, these results indicate significant decreases in the amount and average quality of cristae across aging in murine BAT (Figure 6D).

**Figure 6:**
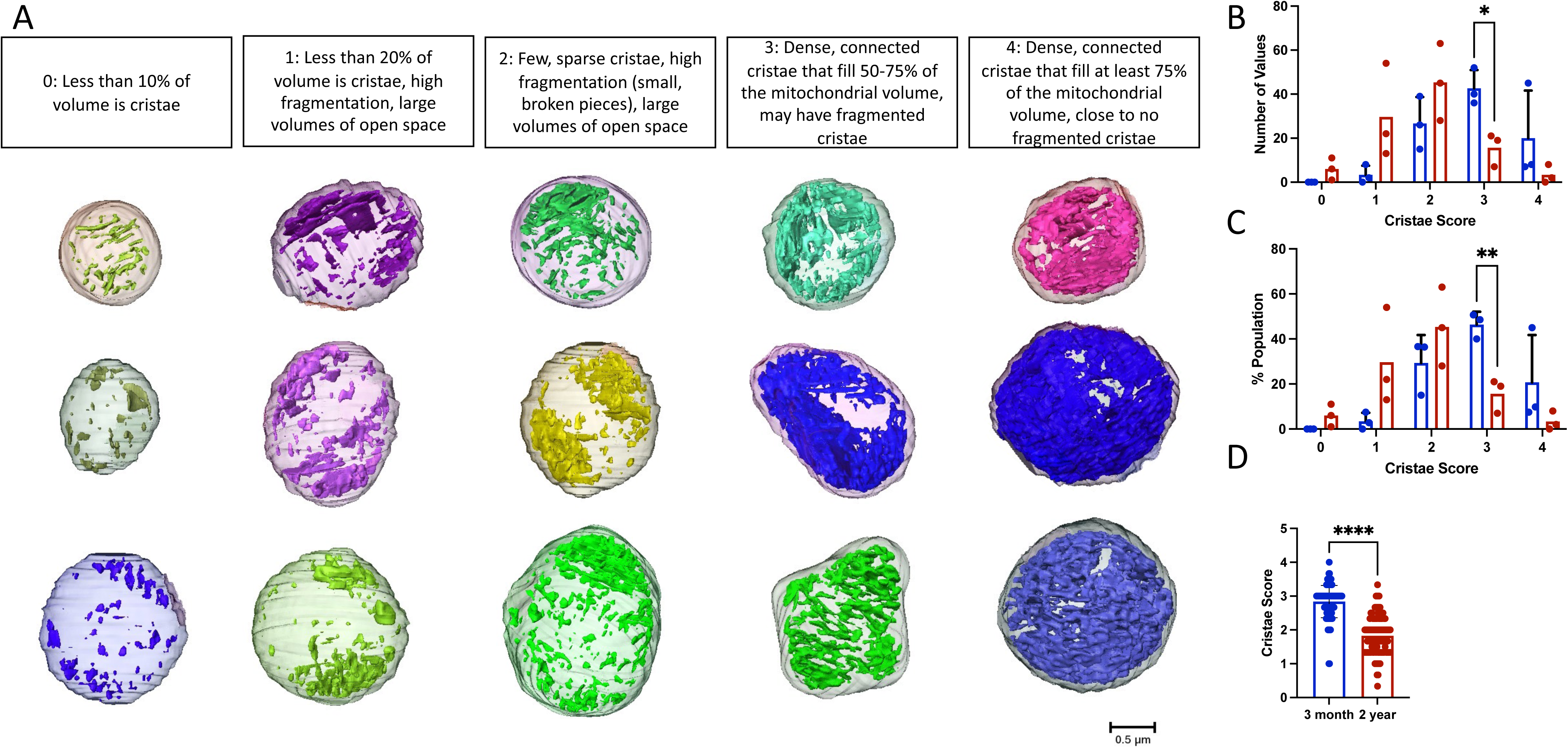
Cristae Score Changes across aging. (A) 3D representative images of mitochondria, from both 3-month and 2-year murine BAT, showing standard cristae morphology for each cristae score. (B) Histogram showing the frequency and (C) percentage of cristae of each score in 3-month and 2-year samples. (D) Comparison of average cristae score in 3-month and 2-year mice. Unpaired t-test was used to evaluate changes, with a variable sample number (n approximately 300), with each mitochondrion represented by a black dot. p < 0.0001 indicated by ****.

### Aged BAT Mitochondria Show No Alterations in Cristae Complexity

Perturbed cristae morphology has been associated with enlarged and dysfunctional mitochondria ^[7, 38]^. Observations of aberrant cristae in senescent cells have also been noted as being partially lost, totally lost, or in a circular formation ^[38]^. To understand the morphological changes that occur with cristae across aging, cristae morphology was examined in the 3-month and 2-year aged samples from a transverse and longitudinal view (SFigure 3A-B). Cristae-otyping, the equivalent to mito-otyping for cristae, was utilized to compare cristae morphology in samples from 3-month and 2-year aged groups across volumes (SFigure 3C). The 2-year aged sample showed the presence of mitochondria with partially lost cristae, along with circular cristae. Notably, we also observed “island” shaped cristae which were smaller and fragmented. The findings indicate that cristae in 2-year samples are more heterogeneous in size and generally smaller in 3-month samples when normalized by mitochondrial volume. Furthermore, they have reduced quality as shown by reductions in cristae score and changes in distribution illustrated by cristae-otyping.

## DISCUSSION

In this study, we examined the mitochondrial ultrastructure changes that occur in murine BAT with aging using SBF-SEM and 3D reconstruction. With the utilization of these tools, increases in mitochondrial 3D volume, perimeter, 3D area, and complexity index as well as decreases in sphericity were quantified between adult and aged murine BAT. Cristae from adult and aged murine BAT showed significant decreases in 3D volume, perimeter, 3D area, and cristae score when compared to the young samples (Figures 4-6). Together, these findings highlight the importance of considering changes in overall cristae density and not relying solely on changes in mitochondrial mass to quantify mitochondrial morphology changes. Although mitochondrial mass changes provide indications of mitochondrial health, cristae, which drive oxidative phosphorylation, should be considered as a method of examining functional quality alterations as well. The results presented here also indicate that both mitochondrial and matrix volumes increase with aging in BAT. Notably, studies have shown that mitochondrial swelling occurs with mitochondrial dysfunction and membrane potential loss ^[38–40]^. Under extreme conditions, osmotic swelling of the mitochondrial matrix follows the opening of the inner membrane permeability transition pore (PTP) and can also irreversibly engage in apoptosis ^[41–43]^.

An interesting observation in our study was the presence of circular cristae in the aged BAT samples that were not present in the young BAT samples. Circular cristae have been described in senescent cells by others and differ from onion cristae ^[44]^ given the lack of multiple rings of cristae present ^[38]^. However, a lack of 3D analysis of the cristae morphology from these studies makes it difficult to compare cristae phenotypes. Beyond this, it is evident that the aging process increases overall mitochondrial cristae density, although the underlying molecular causes have yet to be elucidated (Figure 5C).

An important limitation to this study that should be mentioned is that only male mice were used. Notably, human BAT demonstrates a high expression of estrogen receptors, with activity supplemented by the expression of estrogens and androgens ^[45]^. It is well understood that across aging, estrogen levels are lost which negatively impacts mitochondrial function ^[46]^. Interestingly, it has been found that estrogen can restore dysfunctional mitochondrial phenotypes, reducing the amount of donut-shaped mitochondria in aged monkey brain samples ^[47]^. Past studies looking at female models of mitochondria structure in BAT show that at baseline females have higher expression of UCP1, cristae density, and resilience to stress conditions ^[48]^. Furthermore, in humans, across aging, there is a smaller loss of BAT mass in females ^[25]^. Therefore, future studies are needed to investigate if sex-dependent differences in BAT mitochondrial structure across aging exist and elucidate the role, if any, estrogen-mediated mitochondrial structure protection plays in these differences.

Although the findings here show clear mitochondrial morphology changes with aging, it should also be noted that murine and human BAT have several key differences. For example, while certain studies have demonstrated that β3-adrenergic receptor (β3AR) agonists show protection against diet-related obesity through stimulation of BAT in murine models, this has not been the case for humans ^[49]^. While this study examined interscapular BAT, which is present throughout murine lifespan, it is only present in human babies ^[50]^. One avenue to explore would be looking at specifically more abundant natal-related BAT in human mitochondria compared to the much more limited BAT observed in aged humans to observe if there may be differences in phenotypes that aid to explain the rapid loss of BAT across aging in humans ^[32, 51]^. Therefore, future experiments may consider looking at how mitochondria may present different structures in human models of BAT which may elucidate some of the *in vitro* differences between murine and humans. Similarly, studies have shown that while cristae architecture remains pertinent for energetic capacity across models, aging shows differed functional and structural changes in mice and *Drosophila* ^[52]^. Further studies are needed to understand morphological changes of mitochondria and cristae in human BAT in comparison to murine BAT. Another important limitation of this study that should be noted is that other studies have shown distinct populations of adipocytes, with higher-thermogenic activity populations, which can dynamically form in exposure to cold ^[53]^. Our study did not account for this heterogeneity in the population, which may account for some of the variation observed in mitochondrial structure. Future studies may consider 3D tissue profiling to see if mitochondrial and cristae structure varies across populations of adipocytes or alterations in gene expression ^[53]^.

In summary, to our knowledge, we are the first to elucidate the alterations in 3D mitochondrial and cristae structures across the murine aging process in BAT. Of relevance, this aids to establish standards for the phenotypes presented. Importantly, the phenotypes found here may be related to pre-stress states, alterations in membrane potential, changes in fusion or cristae proteins, sex-dependent differences, or changes in metabolic pathways but further studies are needed to confirm or exclude these possibilities. One avenue that may be altered, in addition to mitochondrial complexity, is the juxtaposition of mitochondria with other organelles which may also uniquely modulate mitochondrial function ^[54]^. Although this study did not delve into the molecular changes that influence the aging phenotypes observed, by establishing these 3D structures, future studies investigating the influence of aging in BAT on all previously mentioned factors can better understand how these factors may relate to specific phenotypes. In the future, this may aid in establishing standards regarding mitochondria and cristae structures, to aid research aimed at developing interventions to mitigate dysfunction during aging in BAT.

## Acknowledgments

Funding by the UNCF/Bristol-Myers Squibb E.E. Just Faculty Fund, BWF Career Awards at the Scientific Interface Award, BWF Ad-hoc Award, NIH Small Research Pilot Subaward to 5R25HL106365-12 from the National Institutes of Health PRIDE Program, DK020593, Vanderbilt Diabetes and Research Training Center for DRTC Alzheimer’s Disease Pilot & Feasibility Program. T32, number DK007563 entitled Multidisciplinary Training in Molecular Endocrinology to A.C. and J.E.S.; T32, number DK007563 entitled Multidisciplinary Training in Molecular Endocrinology to Z.V. CZI Science Diversity Leadership grant number 2022-253529 from the Chan Zuckerberg Initiative DAF, an advised fund of Silicon Valley Community Foundation (to A.H.J). NIH R01DK132669 (to E.D.D). NSF EES2112556, NSF EES1817282, NSF MCB1955975, and CZI Science Diversity Leadership grant number 2022-253614 from the Chan Zuckerberg Initiative DAF, an advised fund of Silicon Valley Community Foundation (to S.D.). NSF grant MCB #2011577I to S.A.M.

---

**Supplemental Figure 1:**
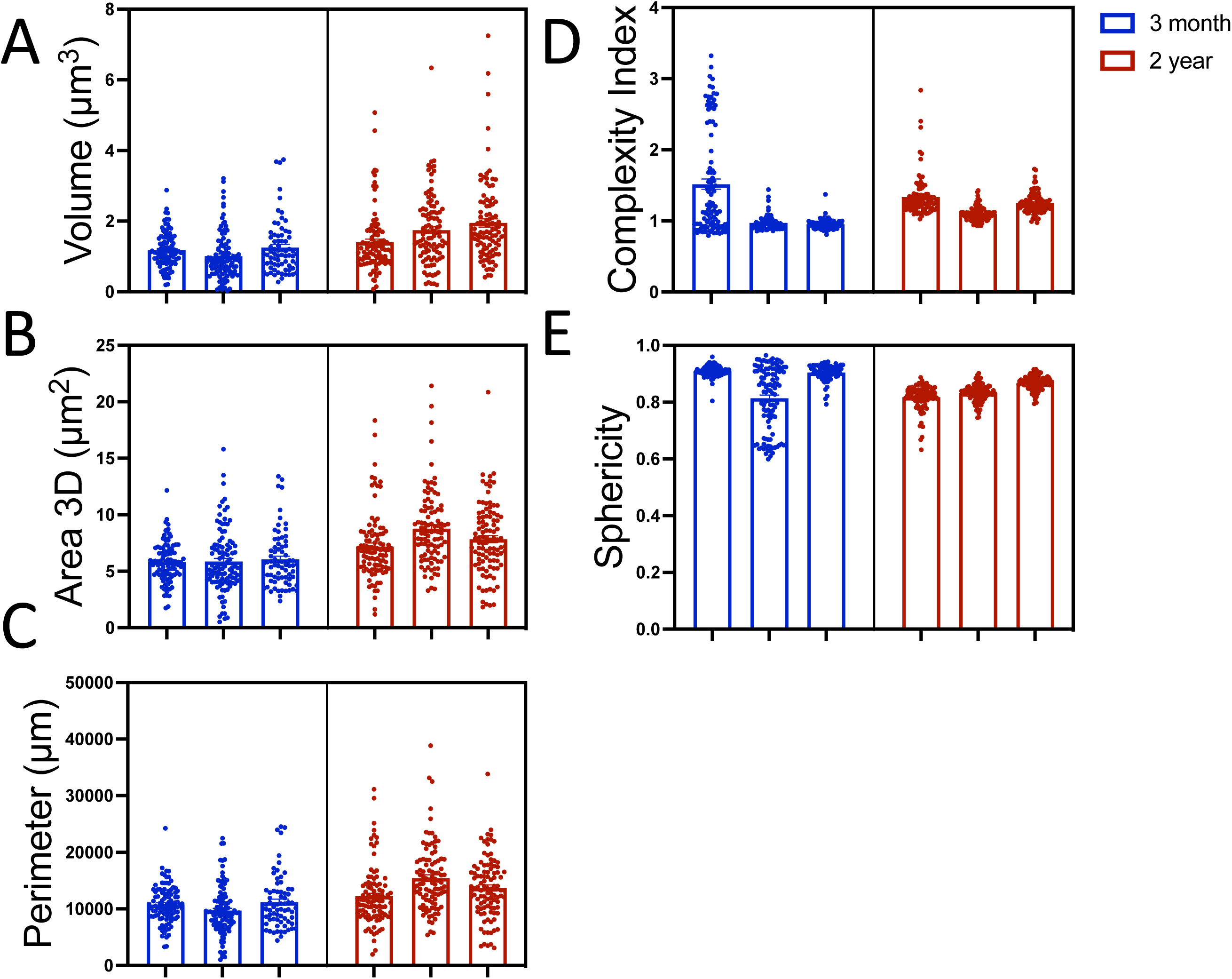
Display of raw mitochondrial quantifications. Each bar represents one of the three mice surveyed at respective age points while dots represent individual mitochondria (n is approximately 300). Values are displayed for (A) mitochondrial volume, (B) area, (C) perimeter, (D) complexity index, and (E) sphericity.

**Supplemental Figure 2:**
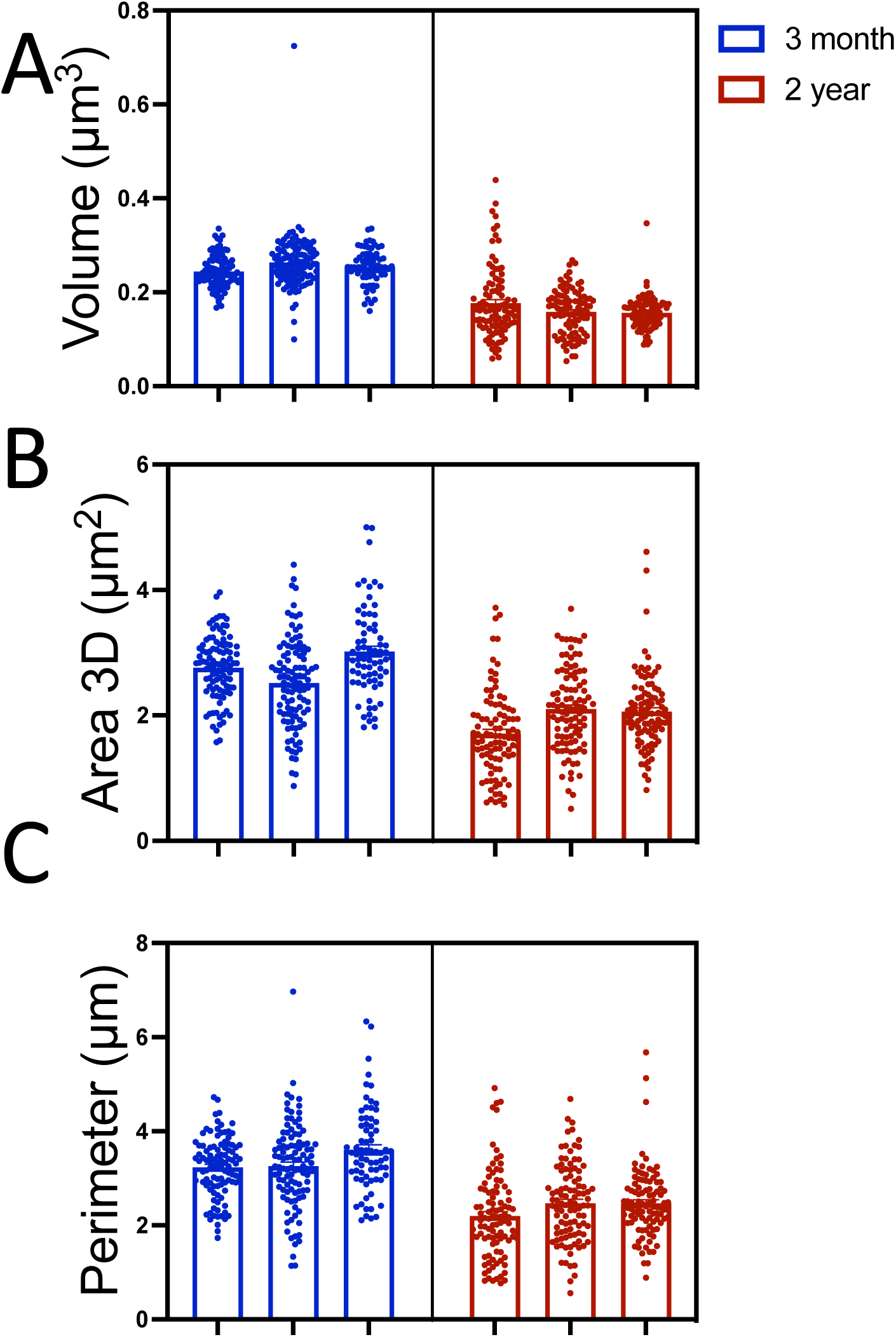
Display of raw cristae quantifications. Each bar represents one of the three mice surveyed at respective age points while dots represent individual mitochondria (n is approximately 300). Values are displayed for (A) cristae volume, (B) area, and (C) perimeter.

**Supplemental Figure 3:**
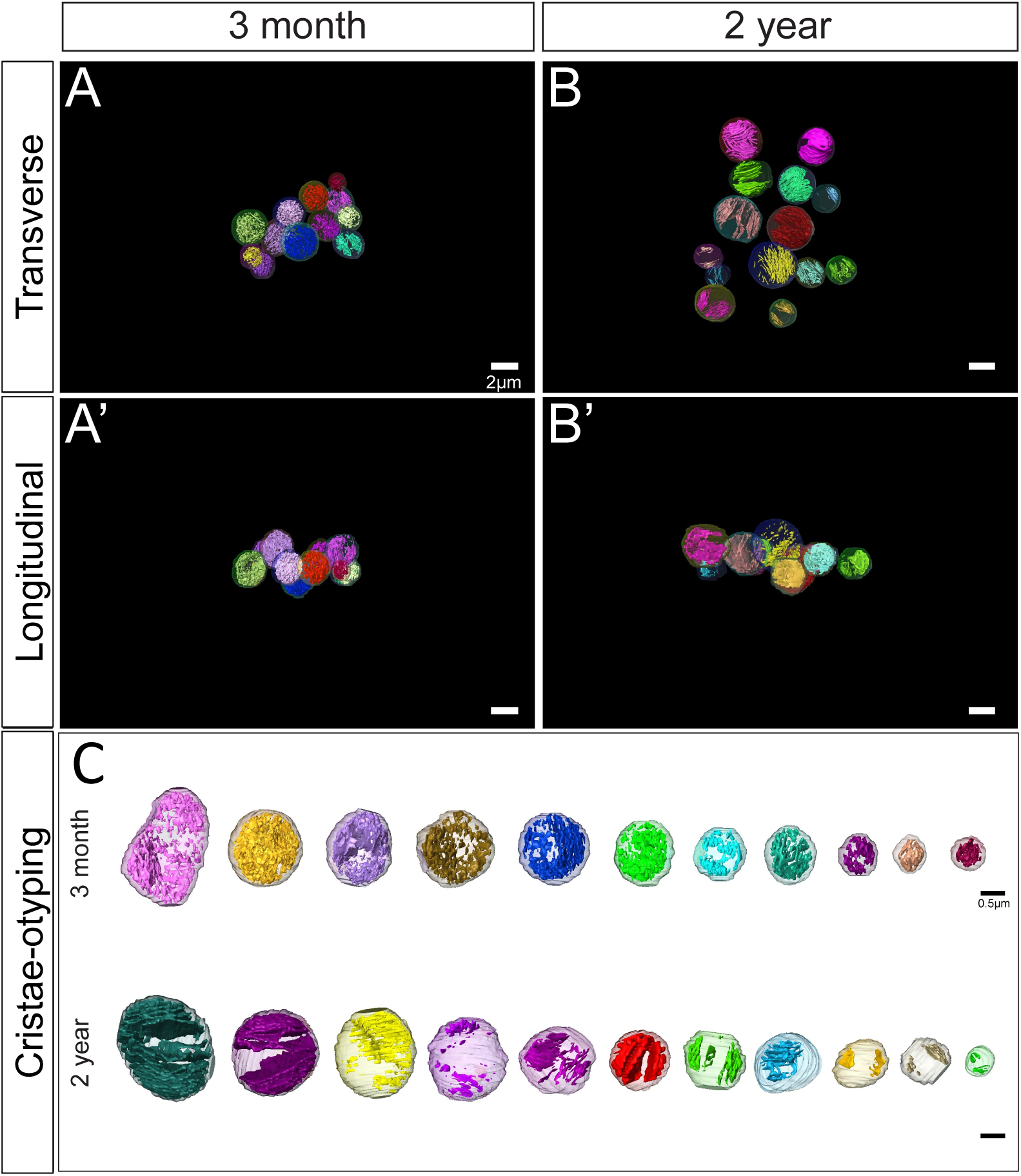
Overall cristae-otyping changes across aging. (A) 3-month murine BAT 3D reconstruction of cristae viewed from the transverse and (A’) 3D reconstruction viewed from a longitudinal point of view. (B) 2-year murine BAT 3D reconstruction of cristae viewed from the transverse point of view (B’) 3D reconstruction viewed from a longitudinal point of view. (C) Cristae-otyping to compare cristae morphology in samples from 3-month and 2-year across volumes.

